# Expression of *ACE2*, the SARS-CoV-2 receptor, and *TMPRSS2* in prostate epithelial cells

**DOI:** 10.1101/2020.04.24.056259

**Authors:** Hanbing Song, Bobak Seddighzadeh, Matthew R. Cooperberg, Franklin W. Huang

**Author notes:** Corresponding Author: Franklin W. Huang, MD PhD, Campus Box 1346, University of California, San Francisco, 513 Parnassus Avenue, HSE 1426, San Francisco, CA 94143-0570.

## Abstract

The COVID-19 pandemic has spread across more than 200 countries and resulted in over 170,000 deaths. For unclear reasons, higher mortality rates from COVID-19 have been reported in men compared to women. While the SARS-CoV-2 receptor *ACE2* and serine protease *TMPRSS2* have been detected in lung and other tissues, it is not clear what sex differences may exist. We analyzed a publicly-available normal human prostate single-cell RNA sequencing dataset and found *TMPRSS2* and *ACE2* co-expressing cells in epithelial cells, with a higher proportion in club and hillock cells. Then we investigated datasets of lung epithelial cells and also found club cells co-expressing *TMPRSS2* and *ACE2*. A comparison of *ACE2* expression in lung tissue between males and females showed higher expression in males and a larger proportion of *ACE2*+ cells in male type II pneumocytes, with preliminary evidence that type II pneumocytes of all lung epithelial cell types showed the highest expression of *ACE2*. These results raise the possibility that sex differences in *ACE2* expression and the presence of double-positive cells in the prostate may contribute to the observed disparities of COVID-19.

## Introduction

As of April 23th, 2020, the COVID-19 pandemic has infected more than 2.5 million people across 213 countries and killed over 170,000 [1]. Despite there being approximately equal number of cases between men and women, emerging reports across countries indicate a higher mortality from SARS-CoV-2 in men than women, though the underlying reasons remain unclear [2]. The extent to which this disparity is due to biological rather than behavioral or comorbid sex differences is unknown.

The SARS-CoV-2 receptor, *ACE2*, and serine protease, *TMPRSS2*, are expressed in lung and other tissues implicated in the clinical manifestations of COVID-19. However, less is known about the exact cell types expressing *ACE2* and *TMPRSS2* that serve as cells of entry and pathogenesis for SARS-CoV-2 [3]. Intriguingly, *TMPRSS2*, which is one of the most dysregulated genes in prostate cancer, is highly expressed in human prostate epithelial cells and is androgen-responsive [4]. The presence of *TMPRSS2* in the human prostate and its regulation by androgen raises the possibility that the prostate may be susceptible to SARS-CoV-2 infection, and that male-sex may be a biological risk factor.

Given the high expression of *TMPRSS2* expression in the prostate, we investigated whether *TMPRSS2* and *ACE2* are co-expressed in human prostate epithelial cells, and compared our findings to expression levels found in five lung single-cell datasets. Lastly, we investigate sex differences in the expression of *ACE2* and *TMPRSS2* in lung epithelia.

## Results

Using a publicly-available single-cell RNA sequencing dataset, we analyzed 24,519 epithelial cells from normal human prostates for expression of *ACE2* and *TMPRSS2* [5]. In this dataset, 0.32% of all epithelial cells (78 of 24,519), 0.47% of all stromal cells (10 of 2,113), 0.06% of endothelial cells (1 of 1,586) and 0.22% of leukocytes (1 of 459) expressed *ACE2*. 18.65% of all epithelial cells (4,573 of 24,519), 41.74% of all stromal cells (882 of 2,113), 16.71% of endothelial cells (265 of 1,586) and 52.07% of leukocytes (239 of 459) expressed *TMPRSS2.* Moreover, we found 30 cells that co-expressed *ACE2* and *TMPRSS2* (0.11% of epithelial cells, 0.10% of all cells) from numerous cell types in the prostate: 0.09% of stromal cells (2 of 2,113), 0.06% of endothelial cells (1 of 1,586), 0.03% of basal epithelial cells (5 of 18,439), 0.18% of luminal epithelial cells (4 of 2,238), 0.40% of hillock cells (10 of 2,530), and 0.61% of club cells (8 of 1,312) (Figure 1).

**Figure 1.**
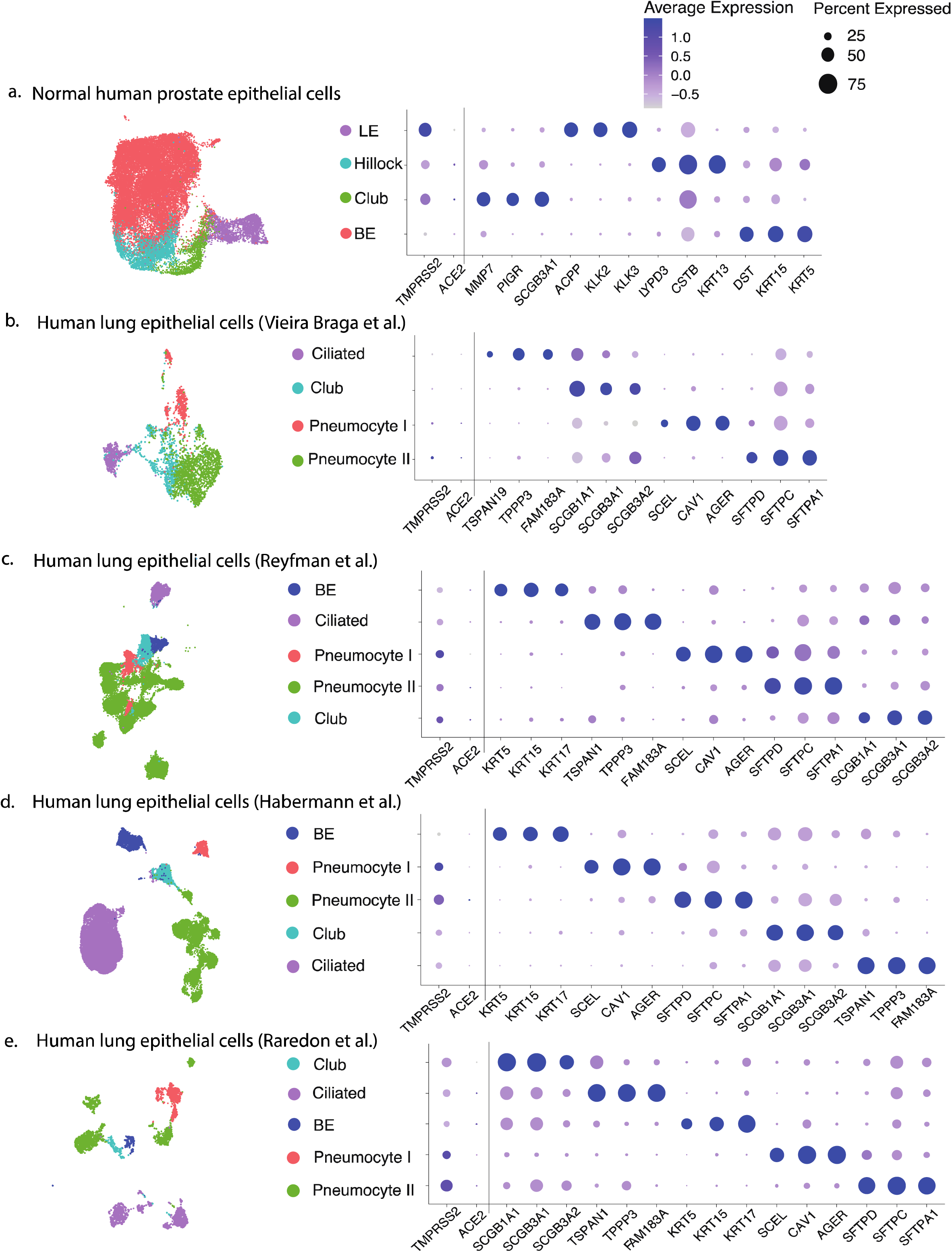
Cell type distribution and top differentially expressed gene marker expression of the four datasets used in the current study. a. Normal human prostate epithelial cells [5] (BE: basal epithelial cells, LE: luminal epithelial cells). b. Human lung epithelial cells from the Vieira Braga et al. study [7]. c. Human lung epithelial cells from the Reyfman et al. study [8]. d. Human lung epithelial cells from the Habermann study [9]. e. Human lung epithelial cells from Raredon study [10]. Each dataset was re-clustered and annotated by different cell types, for which distribution was shown in the Uniform Manifold Approximation and Projection (UMAP). For each dataset, a dot plot was generated showing the percentage of expression and average expression level for the most differentially expressed genes in each cell type as well as *ACE2* and *TMPRSS2*. The marker radius represents the percentage of expression and the color gradient represents the average expression level.

Prostate club cells were found to have the greatest proportion of double-positive cells in the human prostate, and these cells bear a strong resemblance to lung club cells [5]. To characterize *ACE2* and *TMPRSS2* expression in prostate club cells, we compared the expression levels of these genes in lung club cells from one mouse lung [6] and four human lung [7-10] single-cell datasets (Supplementary Table 1). We found a higher proportion of cells expressing *TMPRSS2* than those expressing *ACE2* (Figure 1) in all major lung epithelial cell types (basal epithelial cells, ciliated cells, club cells, type I and II pneumocytes). Specifically, in club cells, we detected the following proportions of double-positive cells in the datasets: 0.33% (7 of 2,113) of human lung club cells in the Reyfman et al. dataset [8]; 0.21% (3 of 1,410) of human lung club cells in the Habermann et al. dataset [9] (Figure 2); and 1.86% (48 of 2,578) of mouse lung club cells in the Montoro et al. dataset [6] (Supplementary Figure 1). Two other human lung datasets showed no such double-positive club cells [7,10] (Figure 2).

**Figure 2.**
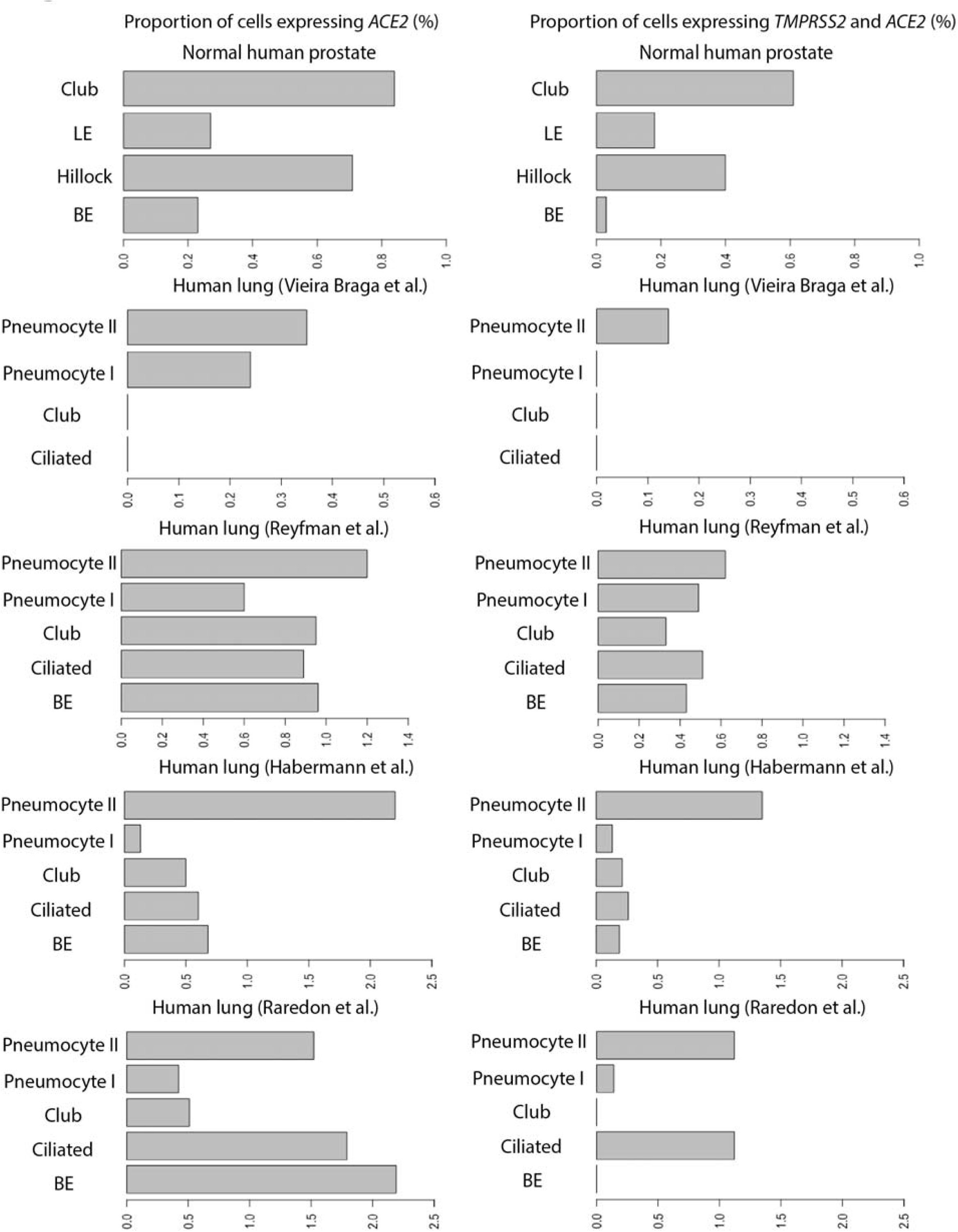
Proportion of cells expressing *ACE2*, and co-expressing of *TMPRSS2* and *ACE2* (%) in each cell type (vertical axes) of the human prostate epithelial cells and four human lung epithelial cell datasets analyzed in the current study.

Lastly, to test sex differences in expression profiles of *ACE2* and *TMPRSS2*, we compared *ACE2* and *TMPRSS2* expression differences within the integrated lung epithelial cells dataset stratified by sex [8,10] (Male: N = 12, Female: N = 19, Supplementary Table 1). Overall, there was no significant difference in *TMPRSS2* expression between males and females in the human lung (Figure 3a). However, *ACE2* expression was higher in males (log 2 normalized expression level: 0.019 in male vs 0.0068 in female, p < 0.001, Figure 3a).

**Figure 3.**
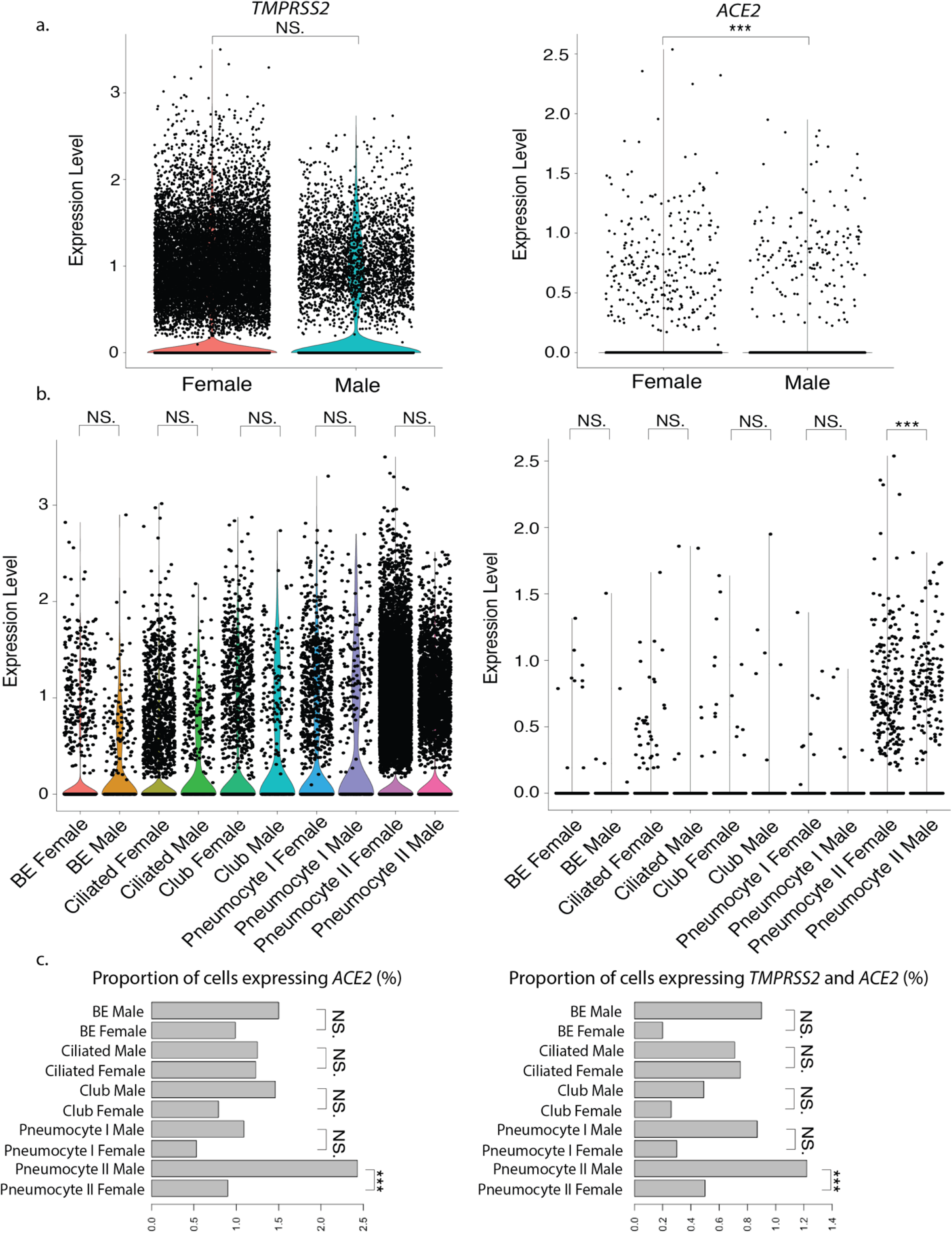
a. Comparison of *TMPRSS2* and *ACE2* expression level between females (N=19) and males (N=12) for all human lung epithelial cells (a) and by each epithelial cell type (b) within the integrated dataset (Supplementary Table 1). Statistical significance levels computed from an unpaired two-samples Wilcoxon test were indicated in the violin plots (NS: non-significant, *: p < 0.05, **: p < 0.01, ***: p < 0.001). c. Proportion of cells expressing *ACE2*, and co-expressing of *TMPRSS2* and *ACE2* (%) in each epithelial cell type and sex of the integrated dataset. Statistical significance levels computed from the Fisher’s exact test were indicated in the bar charts (NS: non-significant, ***: p < 0.001).

To characterize differences in cell types by sex, we then compared *ACE2* and *TMPRSS2* proportions of expression in the integrated dataset. We found higher proportions of type II pneumocytes in males that expressed *ACE2* and co-expressed *ACE2* and *TMPRSS2* compared to type II pneumocytes in females (p < 0.001, Figure 3c). It is not clear if *TMPRSS2* and *ACE2* expression are regulated by the same process but our analysis reveals that their expression levels are positively correlated in lung cell lines (Supplementary Figure 2).

## Discussion

In this study examining *ACE2* and *TMPRSS2* expression in prostate and lung epithelia, we highlight the prostate as a potential organ susceptible to SARS-CoV-2 infection. In human prostate cells, we found hillock and club cells as double-positive cells with the highest proportion of *ACE2* and *TMPRSS2* co-expression. Since both of these proteins are considered to be required for infection, these double-positive cells could potentially serve as reservoirs for SARS-CoV-2 infiltration and damage to the prostate; however, they constitute approximately 0.07% of all prostate epithelial cells. Interestingly, Henry and colleagues recently characterized the prostate club cell [5] as sharing a genetic signature similar to lung club cells [6]. Therefore, in order to better understand the prostate’s susceptibility to infection, we compared *ACE2* and *TMPRSS2* expression in both prostate and lung club cells. We found that 0.61% of prostate club cells co-express *ACE2* and *TMPRSS2*, which correlate with the low numbers of lung club cells with *ACE2 and TMPRSS2* co-expression across four human datasets. Compared to the proportion of *ACE2* and *TMPRSS2* co-expression in type II pneumocytes from prior reports (3.8%) [11], club cells have a markedly lower proportion of double-positive cells. Implications of SARS-CoV-2 infiltration of the prostate via club cells are unclear and warrant further investigation.

Here we also report differences in the co-expression of *ACE2* and *TMPRSS2* in type II pneumocytes by sex. In the integrated dataset with sex data reported, male type II pneumocytes possess a significantly greater proportion of cells expressing *ACE2* and cells co-expressing *ACE2* with *TMPRSS2*. We found no significant difference in *TMPRSS2* expression between male and female lung epithelial cells. This finding is consistent with another recent report examining the same sex difference [12]. If these findings are reproducible with greater sample sizes, the greater proportion of *ACE2* and *TMPRSS2* co-expression in male type II pneumocytes may contribute to the associated disparities in COVID-19 pathogenesis. SARS-CoV-2-related alveolar damage has been implicated to primarily take effect via infiltration of type I and II pneumocytes and ciliated epithelial cells [13]. Type II pneumocytes perform essential functions for alveolar integrity via stem-cell properties and surfactant secretion [14], and their demise can contribute to alveolar collapse and respiratory failure. However, we caution against the generalizability of these findings due to the small sample size, lack of consensus among the data, and the lack of control for confounding variables that may modulate *ACE2* or *TMPRSS2* expression such as smoking [8,10].

Here we find that prostate epithelial cells co-express *TMPRSS2* and *ACE2* with the highest proportion found in hillock and club cells. Whether differences in *TMPRSS2* and *ACE2* expression mediate SARS-CoV-2 pathogenesis and whether androgen signaling can affect COVID-19 disease remain to be studied. Sex differences in *TMPRSS2* and *ACE2* expression could influence potential sites of disease and viral reservoirs. Sex differences in *TMPRSS2* expression alone in the lung may not drive the higher burden of SARS-CoV-2 disease among men. However, our finding that distinct populations of cells in the prostate co-express *TMPRSS2* and *ACE2* raises the possibility that sex differences could influence COVID-19 disease pathogenesis in males differently than females. Further research into *TMPRSS2* expression and its modulation within the lung and other relevant cell types that may impact *ACE2* and SARS-CoV-2 pathogenesis is needed.

## Methods

We adopted previously published single cell RNA-seq datasets of normal human prostate epithelial cells, mouse lung, and four human lung epithelial cells (Supplementary Table 1). Datasets were acquired using the raw count matrices made available by previous studies. For the human prostate, mouse, the Reyfman et al. and the Habermann et al. human lung datasets, cell types were annotated by respective cell type annotation metadata provided in the original studies and validated by top gene markers [5,6,8,9]. The cell type of the epithelial cell population in the Vieira Braga et al., dataset [7] was annotated by the top differentially expressed markers and validated by another previous study [15]. Similarly, the cell type of epithelial cell population in the Raredon et al. dataset [10] was annotated. Sample sex data were acquired for the Reyfman study and the Raredon study [8,10] but unavailable for the other human lung datasets [7,9]. All datasets were analyzed in R using the Seurat 3.1.4 package for visualization and comparison [16]. Gene expression in each dataset was normalized and scaled following the standard Seurat workflow, and sample batch effects were removed using the integration function implemented in Seurat 3.1.4 [16]. Cell distribution in each dataset was visualized in Uniform Manifold Approximation and Projection (UMAP), generated based on the principal component analysis and color-coded by different cell types annotation from the original study. The Reyfman et al. dataset and the Raredon et al. dataset were integrated following the Seurat 3.1.4 integration workflow to remove the batch effect from different samples and stratified by sex of samples reported in the original studies. Top differentially expressed gene markers were identified using the FindAllMarker function built within Seurat and ranked by the log2 fold change of expression [16]. The statistical significance of expression levels between males and females was computed from an unpaired two-samples Wilcoxon test and the proportions of expression between males and females were compared using Fisher’s Exact tests.

## Acknowledgements

This work was funded in part by the Department of Defense W81XWH-17-PCRP-HD (F.W.H., H.S), the National Institutes of Health/National Cancer Institute P20 CA233255-01 (F.W.H., H.S), U19 CA214253 (F.W.H., H.S), and the Prostate Cancer Foundation (F.W.H.).

The authors have declared that no conflict of interest exists.

---

**Supplementary Figure 1.**
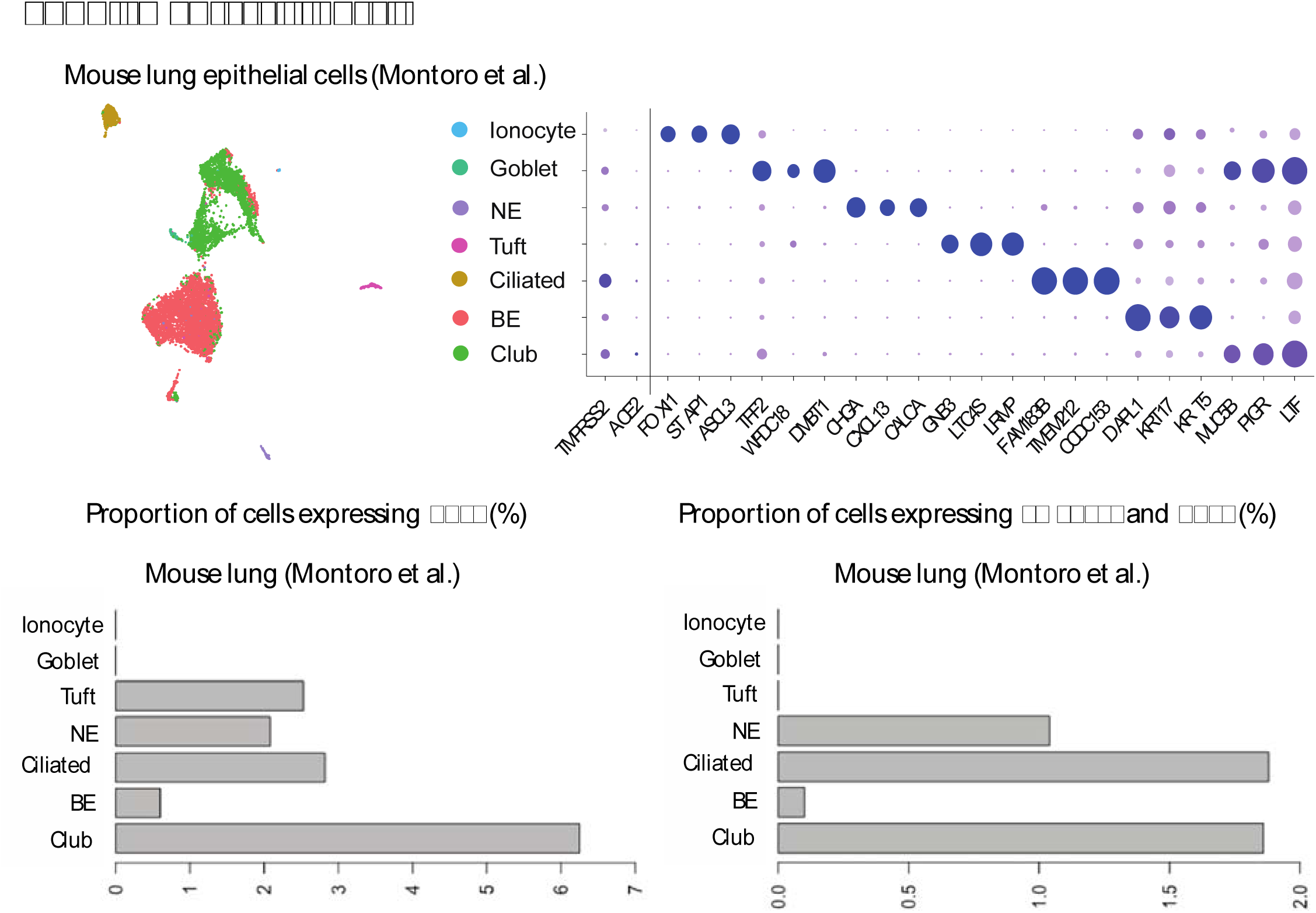
Mouse lung epithelial cells distribution and the proportions of cells expressing *ACE2* and co-expressing of *TMPRSS2* and *ACE2* (%) from the Montoro study [6]. Each dataset was re-clustered and annotated by different cell types, for which distribution was shown in the Uniform Manifold Approximation and Projection (UMAP). A dot plot was generated showing the percentage of expression and average expression level for the most differentially expressed genes in each cell type as well as *ACE2* and *TMPRSS2*. The marker radius represents the percentage of expression and the color gradient represents the average expression level.

**Supplementary Figure 2.**
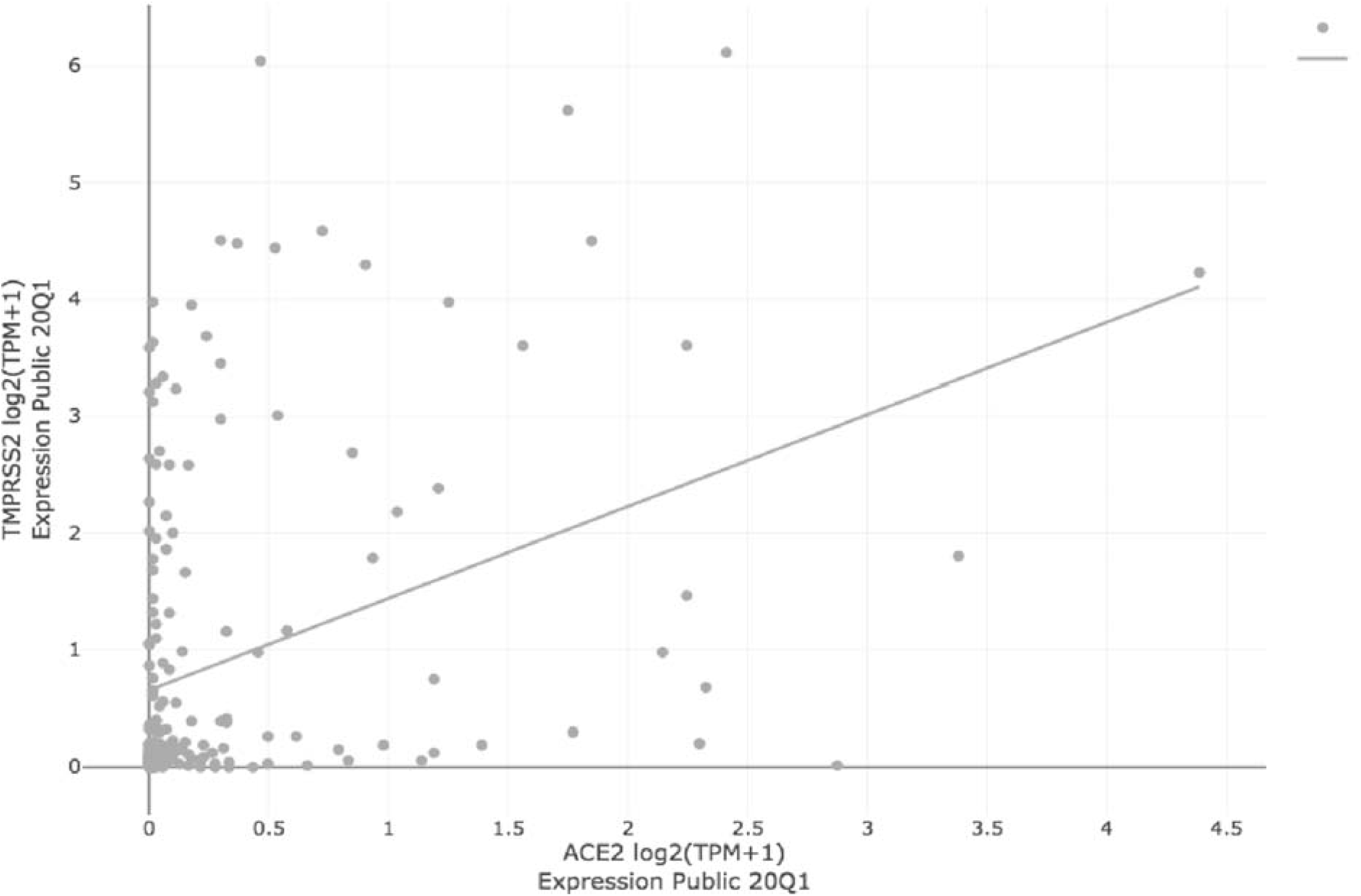
Comparison of *TMPRSS2* and *ACE2* gene expression levels in human lung cancer cell lines (n=206) from the depmap.org portal; Pearson’s correlation = 0.363; p = 8.16 × 10^−8^ (linear regression).

