## Supplementary Material for "Expression of *ACE2*, the SARS-CoV-2 receptor, and *TMPRSS2* in prostate epithelial cells"

**Supplementary Table 1**

| Tissue | Dataset | Species | Number of cells | Source |
| --- | --- | --- | --- | --- |
| Prostate | Henry et al. [5] | Human | 28.702 | Count matrix from GSE117403 |
| Lung | Montoro et al. [6] | Mouse | 7,193 | Count matrix from GSE103354 |
| Lung | Braga et al. [7] | Human | 2,973 | Count matrix from GSE130148 |
| Lung | Reyfman et al. [8] | Human | 34,989 | Count matrix from GSE122960 |
| Lung | Habermann et al. [9] | Human | 28,221 | Count matrix from GSE135893 |
| Lung | Raredon et al. [10] | Human | 4,214 | Count matrix from GSE133747 |
| Lung | Integrated dataset | Human | 39,203 | Integration of the Reyfman and the Raredon datasets [8,10] |
